# A regenerative stem cell-derived matrix accelerates functional dermal wound repair in a preclinical diabetic model

**DOI:** 10.64898/2026.02.20.707094

**Authors:** Colette A. Abbey, Joshua Benton, Erin Goebel, Jocelyn Ma, Sebastian Lomeli, Indu Kancharla, Ivan Juarez, Akshaya Kannan, Christopher Story, Andrew Haskell, Hussain Alcassab, Kayla J. Bayless, Carl Gregory

**Author notes:** **Corresponding authors:** Kayla Bayless, PhD, Associate Professor, Department of Medical Physiology and Institute for Regenerative Medicine, School of Medicine, Texas A&M University, 1114 TAMU | 8447 John Sharp Pkwy, Rm 4412, Bryan, TX 77807, Carl Gregory, PhD, Associate Professor, Department of Medical Physiology and Institute for Regenerative Medicine, School of Medicine, Texas A&M University, 1114 TAMU | 8447 John Sharp Pkwy, Rm 4414, Bryan, TX 77807.

## Abstract

Despite the growing prevalence of non-healing diabetic wounds, no current treatment options overcome multifactorial deficits in repair. To this end, a mesenchymal stromal cell-derived regenerative extracellular matrix (rECM) was evaluated for the ability to accelerate cutaneous wound repair in leptin receptor-deficient (db/db) diabetic mice with paired full-thickness dorsal skin defects. A single dose of rECM significantly accelerated wound closure compared with vehicle controls. Also, rECM dose-dependently improved overall histological healing scores and modulated granulation tissue dynamics, with the highest dose promoting rapid resolution of granulation tissue relative to wound area. Spatial transcriptomics and immunofluorescence revealed that rECM drove robust formation of de novo peripheral nerve clusters characterized by the Schwann cell marker, p75. The rECM also enhanced vascular maturation in healed wounds, increasing average blood vessel size, smooth muscle actin–positive vessels, and vessel density within myofibroblast-rich regions. In a complementary 3D angiogenic sprouting model, rECM accelerated endothelial invasion and filopodia extension, and at higher concentrations induced contraction of collagen matrices consistent with accelerated resolution of granulation tissue. These data demonstrate that rECM accelerates closure of diabetic skin defects by coordinating faster granulation tissue remodeling with enhanced peripheral nerve formation and vascular maturation.

## Introduction

Chronic wounds affect more than 10 million people in the US, with the prevalence rapidly increasing due to aging, diabetes, and obesity.(1–3) In ideal conditions, wound repair is restorative and replenishes healthy skin, but in chronic conditions such as diabetes, pathophysiological changes result in stalled healing and persistence of wounds.(4) Key deficits include loss of extracellular matrix (ECM) architecture,(5, 6) aberrant regrowth of appendages,(7) and decreased cell motility.(5, 8) Thus, a multifaceted treatment capable of restoring matrix integrity to regenerate complex tissues is warranted.

The ECM is indispensable for skin repair, serving as a structural scaffold, cell attachment substrate and signaling complex. During repair, the ECM provides structural support which forms a provisional matrix to guide cell adhesion, migration and proliferation.(9) Impaired ECM integrity in diabetic wounds lowers tensile wound strength and impairs proliferation of dermal progenitors, decreasing granulation tissue formation while increasing neuropathy, scarring and the risk of infection.(5, 10, 11) Similar deficits in the deposition of functional ECM are also observed with obesity and aging.(11, 12) The mechanisms underpinning failure of dermal healing in diabetes are complex, but reduced capacity to support ingrowth of microvasculature and peripheral nerves are major etiological contributors.(13) Ideally, failed wound closure and scarring could be prevented by restoring or supplementing native ECM architecture, regrowing skin appendages, and raising wound strength.(3, 14) Unfortunately, no current therapy meets any of these criteria (3, 15) making development of effective multi-instructive treatment options a priority.

Mesenchymal stem cells, also referred to as mesenchymal stromal cells (MSCs), hold significant potential to overcome current limitations in wound repair based on their well-documented ability to promote tissue regeneration.(16, 17) When live MSCs are applied to experimental skin wounds, healing is accelerated;(18) however, live cell therapy is challenging to translate to widespread clinical use due to logistic and regulatory hurdles.(19–22) An alternative strategy leverages the healing properties of MSCs by purifying the regenerative extracellular matrix (**rECM**) they secrete in large quantities and using it in place of live cells.(23) To this end, our group has optimized synthesis, purification and solubilization of rECM secreted by a clonally-derived line of induced human pluripotent stem cell-derived mesenchymal stromal cells (ihMSCs).(24–26) The clonal nature of the ihMSCs mitigates concerns about reproducibility and utilization of rECM and also removes safety concerns associated with delivery of live cells.(26) In prior studies, ihMSC-derived rECM drove regeneration of bone by facilitating progenitor cell retention, improving cell survival, accelerating osteogenesis, and stimulating angiogenesis.(24, 27, 28) Notably, purified rECM is enriched with collagens XII and collagen VI, which were vital for effective bone repair. (24) These collagens are also integral in maintenance of connective and musculoskeletal tissues and abundant in healthy adult skin.(29–36) Based on the osteoregenerative capabilities previously observed in bone, and the involvement of colVI and colXII in wound healing,(37–39) we reasoned that regenerative extracellular matrix (rECM) derived from MSCs would similarly improve wound repair in skin. Herein, we demonstrate that when administered to experimental dermal injuries established in diabetic leptin receptor deficient mice, rECM drove wound closure, increased remodeling of granulation tissue, enhanced maturation of blood vessels and stimulated ingrowth of peripheral nerve structures, showing promise as a strategy to accelerate functional wound repair in a diabetic model.

## Results

To test the ability of rECM to accelerate the healing of skin wounds, two full-thickness 8 mm diameter dermal punches (hereafter referred to as defects) were applied to the dorsal skin of leptin receptor-deficient (*db/db*) diabetic mice. This strain has clinical relevance because it mimics the pathology of type 2 diabetes, including delayed wound repair.(40, 41) One defect received solubilized rECM delivered in collagen type I matrices, and the other defect received collagen I matrix alone (vehicl*e*). An initial concentration of rECM (75 µL of 100 µg/ml rECM) was selected based on a prior report of the osteogenic activity of rECM *in vitro*.(24) Wound area was measured by analysis of captured digital images taken by observers blinded to treatment and in some cases, these values were confirmed by caliper measurements (**Fig1A**). After a 6 day lag phase with no appreciable size changes in either group, defect closure began to occur in both groups between days 7-13 (**Fig1B,C**, **FigS1**). During this 7-13 day period, closure was significantly accelerated in the rECM treated group (**Fig1B,C**, **FigS1**). After day 16, the majority of defects were nearly fully resolved in both groups (**Fig1B**, **FigS1**).

**Figure 1.**
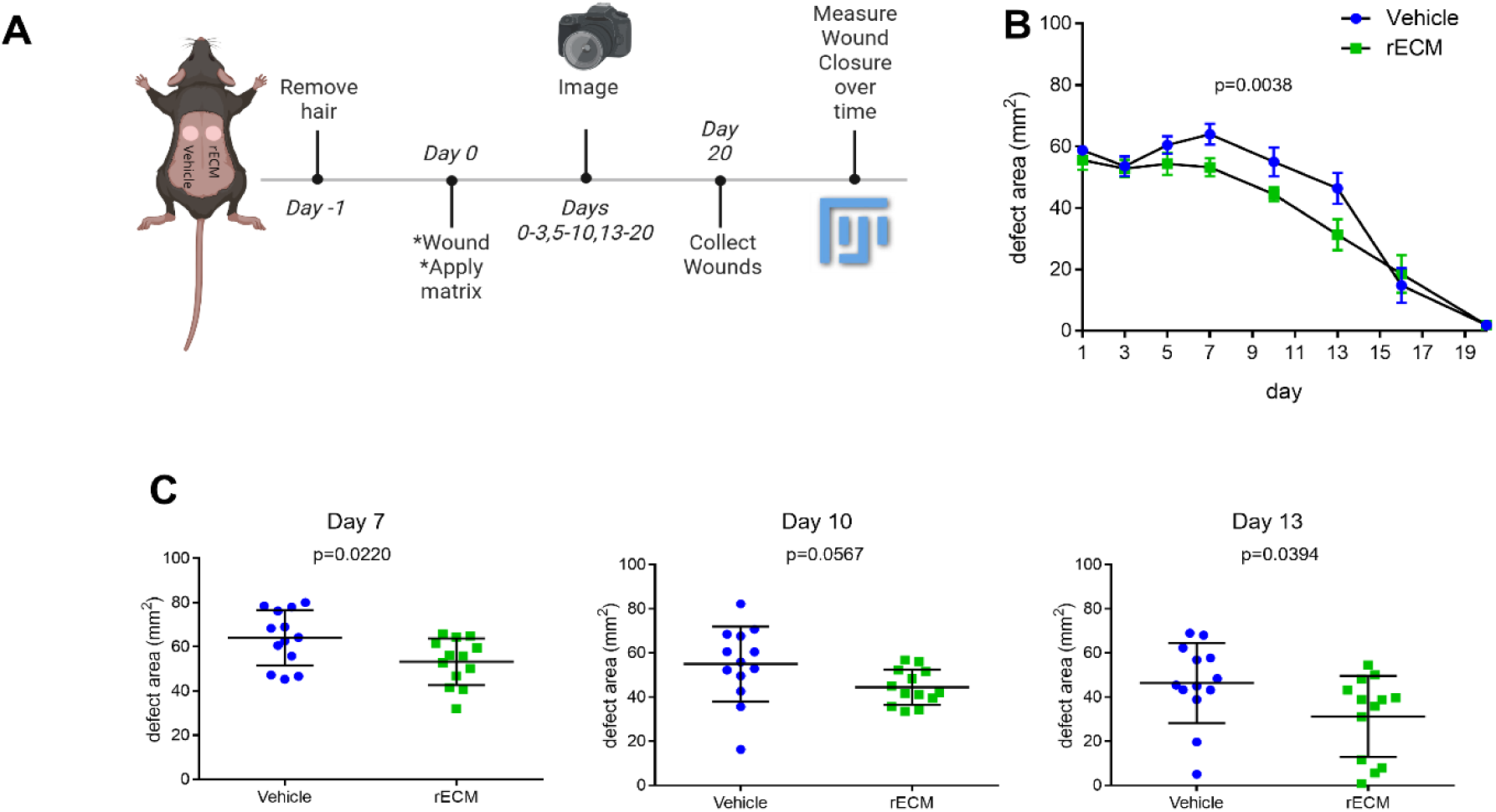
Topical application of rECM accelerates skin wound healing: **Panel A:** Overview of experimental procedures. Two full thickness skin wounds were made on each mouse. Wounds received collagen matrices containing Vehicle or a single dose of rECM. Defect areas were measured by blinded observers using calipers or photographs prior to tissue collection and analysis (Created with Bio-Render.com). **Panel B:** Defect area measurements plotted over time without (Vehicle) or with 100 µg/ml rECM (n = 13-20). **Panel C:** Defect area measurements at Days 7, 10, and 13. *<u>Statistics:</u>* Panel B: 2-way ANOVA; p value represents difference between experimental groups; points indicate means and error bars represent standard error. Panel C: paired t-test; central bars represent mean, and error bars represent standard deviation.

In a second experiment, we tested multiple doses (50, 100, and 200 µg/ml rECM). Analysis of wound area measurements on days 0, 5, 8, and 11 (with doses combined) revealed acceleration of wound closure by rECM (**Fig2A**). Enlargement of the vehicle-treated defects (compared to the calculated area of the 8 mm biopsy punch, 50.26 mm^2^) occurred 2-5 minutes (day 0) after wounding (59.67 mm^2^, SD 10.18 mm^2^, a factor of +19%); however, with administration of rECM to defects, this enlargement did not occur (48.87 mm^2^, SD 11.4 mm^2^ a factor of −2.7%) (**Fig2B**). The observed rapid expansion of the defect area appears to be a genotype-specific phenomenon because a similar observation has been reported by Sullivan *et al.* who compared biopsy punch defects between *db/db* mice and heterozygous controls.(41) Immediately after wounding, *db/db* defects increased by 10% whereas *db/+* controls contracted by 16.7% (when compared to the calculated area of the biopsy punch). In the present study, it is noteworthy that only 2-5 min of exposure to rECM was required to reverse the tendency for enlargement of the defects in *db/db* mice. As in the first experiment, rECM appeared to improve the qualitative appearance of the wounds (**Fig2C**).

**Figure 2.**
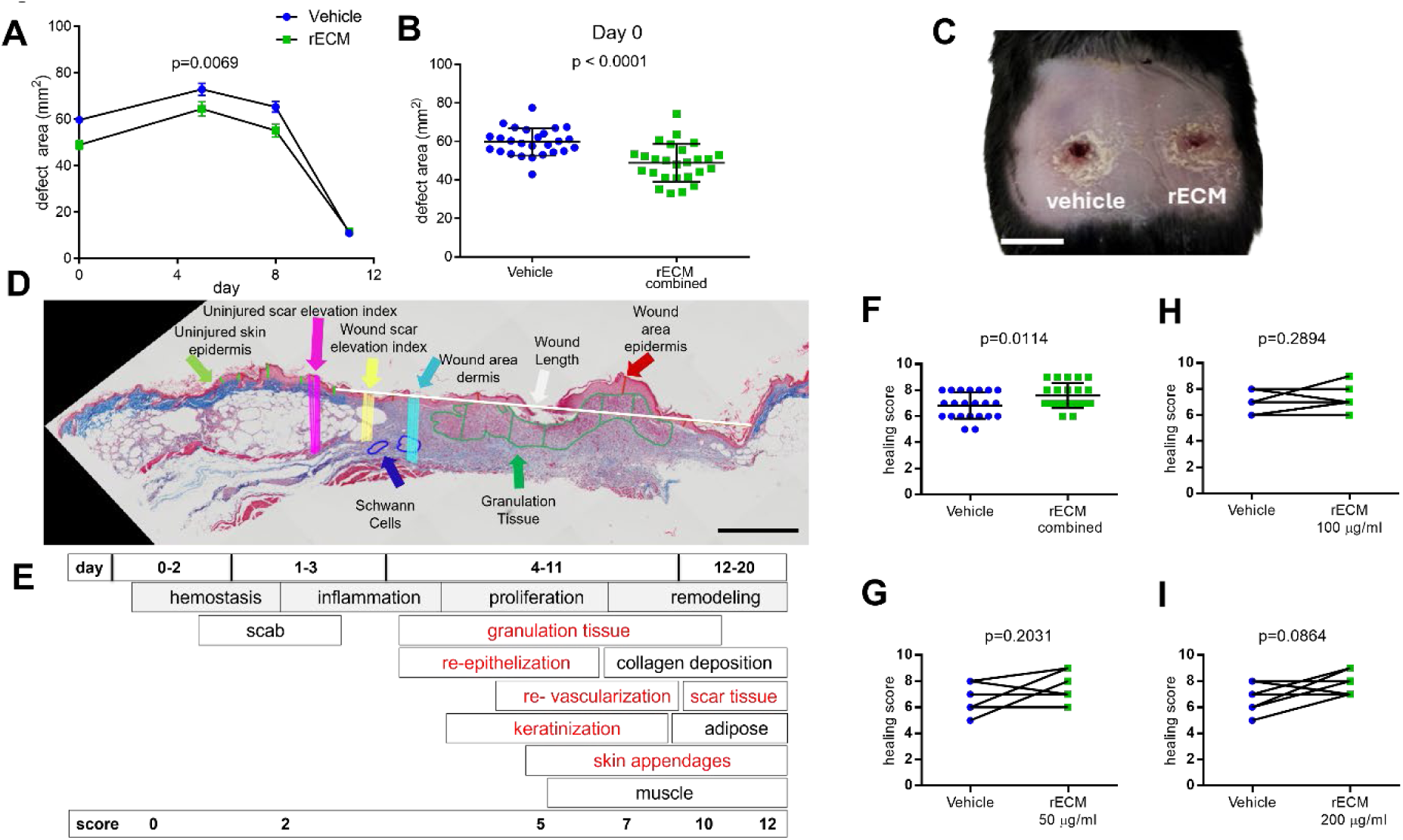
Dose-dependency of rECM in accelerating histological parameters of skin healing: **Panel A:** Defect area measurements for combined rECM doses (50, 100, 200 µg/ml rECM) compared to Vehicle. **Panel B:** Defect area measurements taken immediately after wounding. **Panel C:** Photograph of representative defects at day 11 (scale bar = 10 mm). **Panel D:** Representative Masson’s trichrome-stained wound section with healing parameters used to objectively score skin healing (scale bar = 800µm).**Panel E:** Summary of wound healing phases and completion of key processes over time. Note score increases with completion of each step. The processes listed in red were scored in the study, contributing to a 12-point grading scheme where 0 refers to no healing and 12 refers to completely healed (adapted from van de Vyer *et al.*(42). The detailed scoring system used is outlined in Table 1. **Panel F-I:** Histological Day 11 wound healing scores for Vehicle compared to combined, 50 µg/ml, 100 µg/ml, and 200 µg/ml rECM doses, respectively. *<u>Statistics</u>*: Panel A: 2-way ANOVA; p value represents difference between experimental groups (n=26 mice); points represent means and error bars represent standard error. Panel B: paired t-test; p value represents difference between experimental groups, points represent means and error bars represent standard deviation. Panel F: paired t-test, error bars represent standard deviation and central bar represents mean. Panels G-I: paired t-test, lines connect corresponding control and rECM-treated defects on a single animal (n=6, 7, and 9 for 50, 100, and 200 µg/ml rECM, respectively).

Skin samples were recovered at day 11 for comprehensive histological analysis using an adaptation (**Table 1**) of the methodology of van de Vyer et al.(42) where a healing score is generated based on aggregation of key histological signs of healing including re-epithelialization, epidermal thickness, keratinization, granulation tissue thickness, skin appendage development, and scar elevation index (**Fig2D,E**). At day 11, histological healing scores were significantly higher for the rECM-treated defects when doses were combined and compared to their respective vehicle-treated controls (**Fig2F-I**). The most prominent healing was observed at 200 µg/ml (**Fig2I**).

**Table 1.** Parameters used for histological scoring of wound repair.

| Parameter | Description | Criteria | Score | Adapted Criteria | Score |
| --- | --- | --- | --- | --- | --- |
| Re-epithelization | Complete | 95-100% | 2 | 87-100% | 2 |
|  | Partial | <95%;>0% | 1 | <87%;>0% | 1 |
|  | None | 0% | 0 | 0% | 0 |
| Epidermal thickness index | Normal | 95-105% | 2 | 89-105% | 2 |
|  | Hypertrophy | >105% | 1 | >105% | 1 |
|  | Hypoplasia | <95% | 0 | <89% | 0 |
| Keratinization | Yes |  | 2 | Yes, with epithelium across total wound area | 2 |
|  | No |  | 0 | Yes, with incomplete epithelium across wound area | 1 |
|  |  |  |  | No | 0 |
| Granulation Tissue | Intact dermis | No granular infiltrates | 2 | NA | NA |
|  | Thick GT | >100 µm | 1 | NA | NA |
|  | Thin GT | <100 µm | 0 | NA | NA |
| Remodeling | Complete | Evidence of dermal white adipose, skin appendages, panniculus carnosus | 2 | NA | NA |
|  | Partial | Either/or collagen deposition or dermal white adipose tissue | 1 | NA | NA |
|  | None | No evidence of remodeling | 1 | NA | NA |
| Scar Elevation Index | Normal | 95-105% | 2 | 76-105% | 2 |
|  | Hypertrophied | >105% | 1 | >105% | 1 |
|  | Hypoplasia | <95% | 0 | <76% | 0 |

When analyzing individual parameters, rECM had a positive influence on % re-epithelialization, epidermal thickness and scar elevation index, but these parameters did not attain statistical significance (**Fig3A-C**). On the other hand, with respect to granulation tissue thickness, a biphasic response was observed where 50 μg/ml rECM resulted in an increase in granulation tissue thickness (p=0.0392), and 200 μg/ml resulted in a decrease in granulation thickness (p = 0.0595) compared to controls with the intermediate (100 μg/ml) dose causing no discernible change in granulation tissue thickness (**Fig3D**). Granulation tissue formation typically peaks after the inflammatory stage of healing to drive the initial stages of wound contraction, but it is gradually cleared to facilitate remodeling.(43–45) Therefore, a rapid execution and resolution of the granulation tissue phase is likely to expedite wound closure and transition the final stages of repair. To gain further insight into the potential role of rECM in acceleration of granulation tissue turnover during healing, the proportion of granulation tissue was normalized to wound area. There was no significant change with combined doses (**Fig3E**), but granulation tissue was dose-dependently reduced with rECM treatment, and many defects completely lacked granulation tissue after 11 days in the 200 μg/ml rECM treatment group (**Fig3E**). Together, these data indicate that rECM has the capacity to accelerate skin defect healing through mechanisms involving acceleration of granulation tissue resolution and wound contraction. Histology was performed on day 11 to maximize the probability of observing differences in wound size, angiogenesis and granulation tissue but this time-point was too early to effectively quantify *de novo* skin appendages such as hair follicles.(46) Spatial sequencing of these specimens combined with spot deconvolution analysis using K means clustering facilitated detailed wound characterization in the presence of rECM. As expected, muscle, blood vessel growth, epithelium and keratinocytes, neurogenic tissue and cytokine-rich inflammatory tissue was detectable in both replicates (**FigS2, TableS1**) indicating that rECM did not detrimentally or fundamentally alter the histological and molecular hallmarks of skin repair.

**Figure 3:**
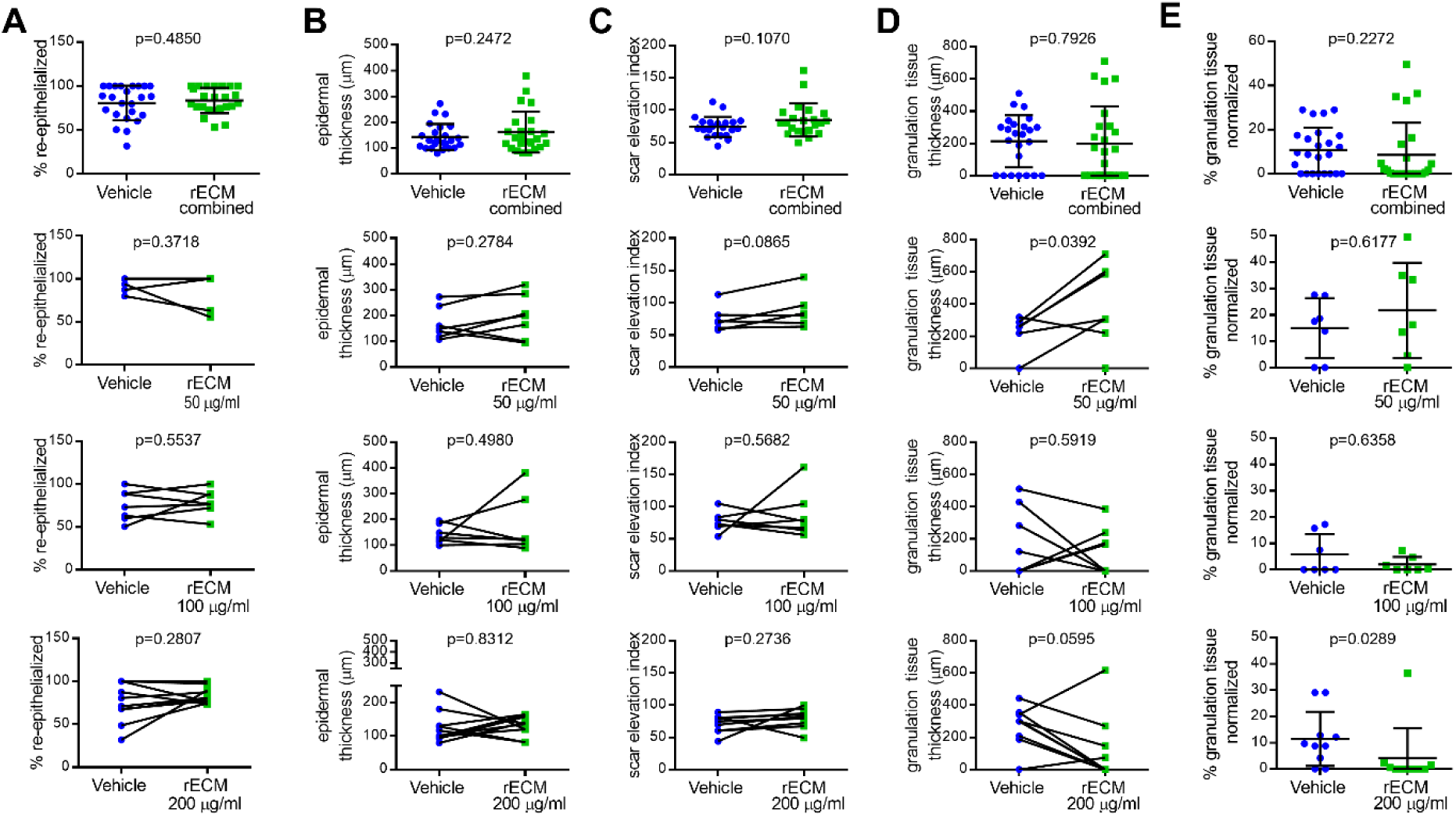
Individual measurements of histological parameters as in Figure 2. **Panel A:** Percent re-epithelialization measurements for combined doses of rECM compared to Vehicle control. From top to bottom: combined doses rECM, 50 µg/ml, 100 µg/ml and 200 µg/ml rECM. **Panel B:** As Panel A, but epidermal thickness measurements. **Panel C:** As Panel A, but scar elevation index. **Panel D:** As Panel A, but granulation tissue thickness. **Panel E:** As Panel A, but percent granulation tissue normalized to defect area. *<u>Statistics</u>*: Panels A-D: For combined doses, paired t-test, n=24; central bar represents mean and error bars represent standard deviation. For individual doses, paired t-test (n=7-10), lines connect corresponding control and rECM-treated defects on a single animal. Panel E: Mann-Whitney test (n=7, 7, and 10 for 50, 100, and 200 µg/ml rECM, respectively).

While the morphology and molecular biology of rECM-driven healing was largely unremarkable, there was a noteworthy prevalence of structures that resembled early-stage peripheral nerve clusters in rECM-treated defects (**Fig4A**). Spatial transcriptomic analysis of these structures (**Fig4B**) revealed the areas of interest exhibited high and exclusive expression of established Schwann cell markers,(47) including Mpz (P0), Gap43, Sox10 and Krox20 (Egr2). A list of upregulated genes in the clusters (**Table S2**) (compared with the entire dataset) was generated (Fig4C) and used to interrogate the GOnet (DICE Tools) (48, 49) and PanglaoDB (50) databases to determine the identity of the structures. The most highly represented features listed by GOnet were ensheathment of neurons, gliogenesis, myelination, axon ensheathment and glial cell differentiation. The list also flagged oligodendrocytes, neurons, astrocytes and Schwann cells in the Pangaloo database. Overall, these data indicated a strong signature that corresponded to Schwann cells and peripheral nerves. Immunofluorescence staining for p75, a marker for Schwann cells, revealed elevated p75 staining in rECM treated wounds, but little staining in controls (**Fig4D,E**). The peripheral nerve clusters had increased area (**Fig4F**) and were more numerous (**Fig4G**) in rECM-treated specimens (with doses combined), with the highest dose of rECM (200 μg/ml) generating the most clusters (**Fig4H**). Collectively, these data indicate that rECM, especially at higher doses, drives growth of *de novo* peripheral nerves in the skin defects in addition to accelerating the resolution of granulation tissue.

**Figure 4.**
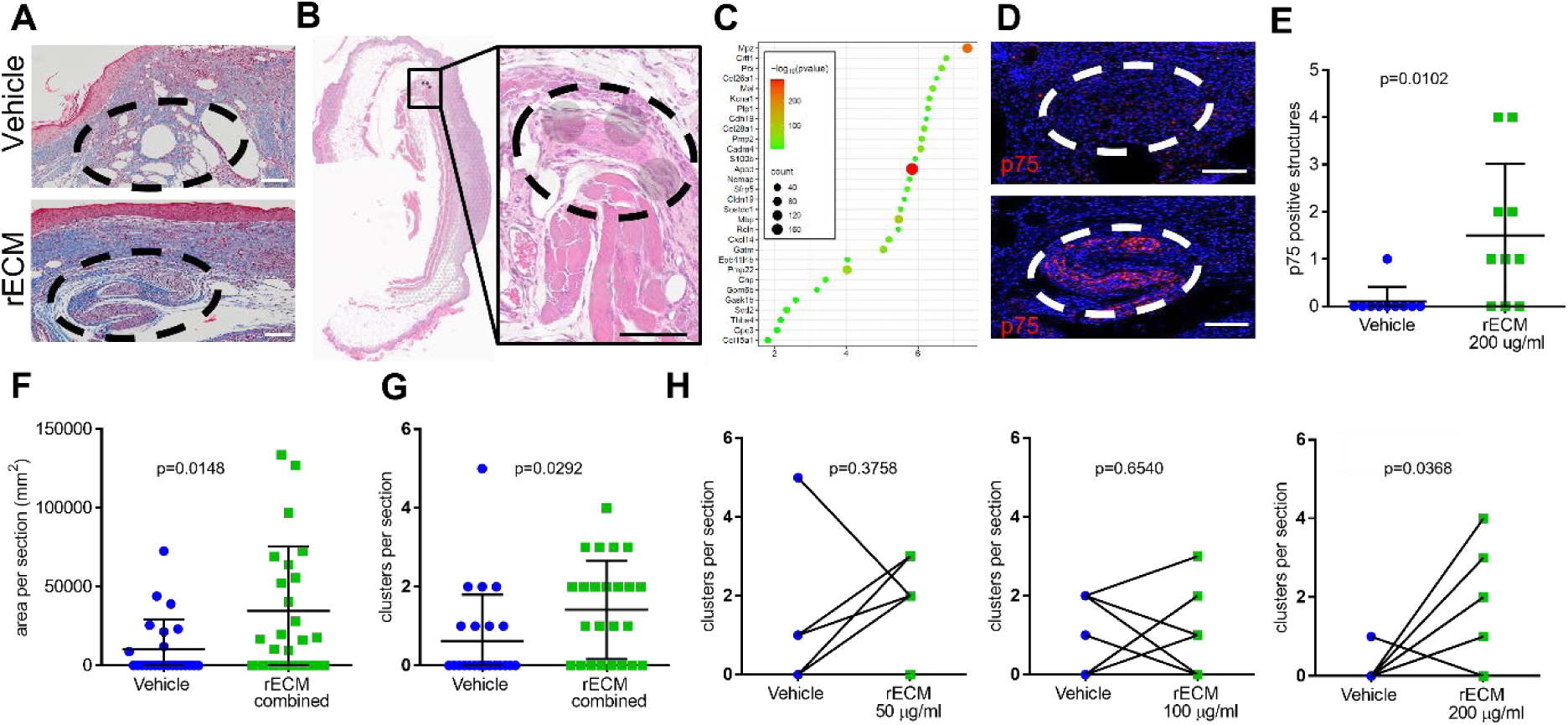
rECM treatment increased the number and size of peripheral Schwann cell clusters in healing skin. **Panel A:** Masson’s Trichrome stained sections illustrate morphology of clusters in healing skin (scale bar = 100 µm). **Panel B:** Spatial sequencing of a healing skin section with circular overlays representing individual transcriptomic datasets. Gray circle represents enrichment for transcriptomic Schwann cell markers (scale bar = 100 µm). **Panel C:** Inset summarizes gene expression frequency from SRPLOT analyses(https://www.bioinformatics.com.cn/plot_basic_gopathway_enrichment_bubbleplot_081_en). **Panel D:** Immunofluorescent staining of p75-positive Schwann cell clusters in healing skin (scale bar = 100 µm). **Panel E:** Number of p75-positive structures counted per histological section for 200 µg/ml dose of rECM. **Panel F:** Area of peripheral nerve cell clusters measured per histological section for combined doses of rECM. **Panel G:** Number of peripheral nerve cell clusters counted per histological section for combined doses of rECM. **Panel H:** As in G but plotted for each dose of rECM. *<u>Statistics:</u>* Panels E, F, and G: paired t-test; central bar represents mean, and error bars represent standard deviation. Panel H: paired t-test, lines connect corresponding control and rECM-treated defects on a single animal; (n=7, 7, and 10 for 50, 100, and 200 µg/ml rECM, respectively).

As nerves and blood vessels typically develop in close proximity and are guided by common cues, (51–53) we next analyzed blood vessel formation and maturation in response to high doses of rECM treatment (200 µg/ml). Immunofluorescent staining of CD31 and smooth muscle actin (SMA) to label endothelial and smooth muscle cells, respectively, revealed areas of robust new blood vessel growth (**Fig5A**). Analysis of immunofluorescent CD31 staining revealed that the average blood vessel size was larger with rECM treatment (**Fig5B**). In addition to increased size of blood vessels, the number of SMA-positive blood vessels was also increased (**Fig5C**). Further, when areas of myofibroblast staining were isolated for analysis (**Fig5D**), the density of blood vessels in close proximity to myofibroblasts was also increased (**Fig5E**). Collectively, these results show that higher doses of rECM accelerate vascular maturation, consistent with accelerating the resolution of granulation tissue and enhancing the growth of peripheral nerves in diabetic mouse skin wounds.

**Figure 5.**
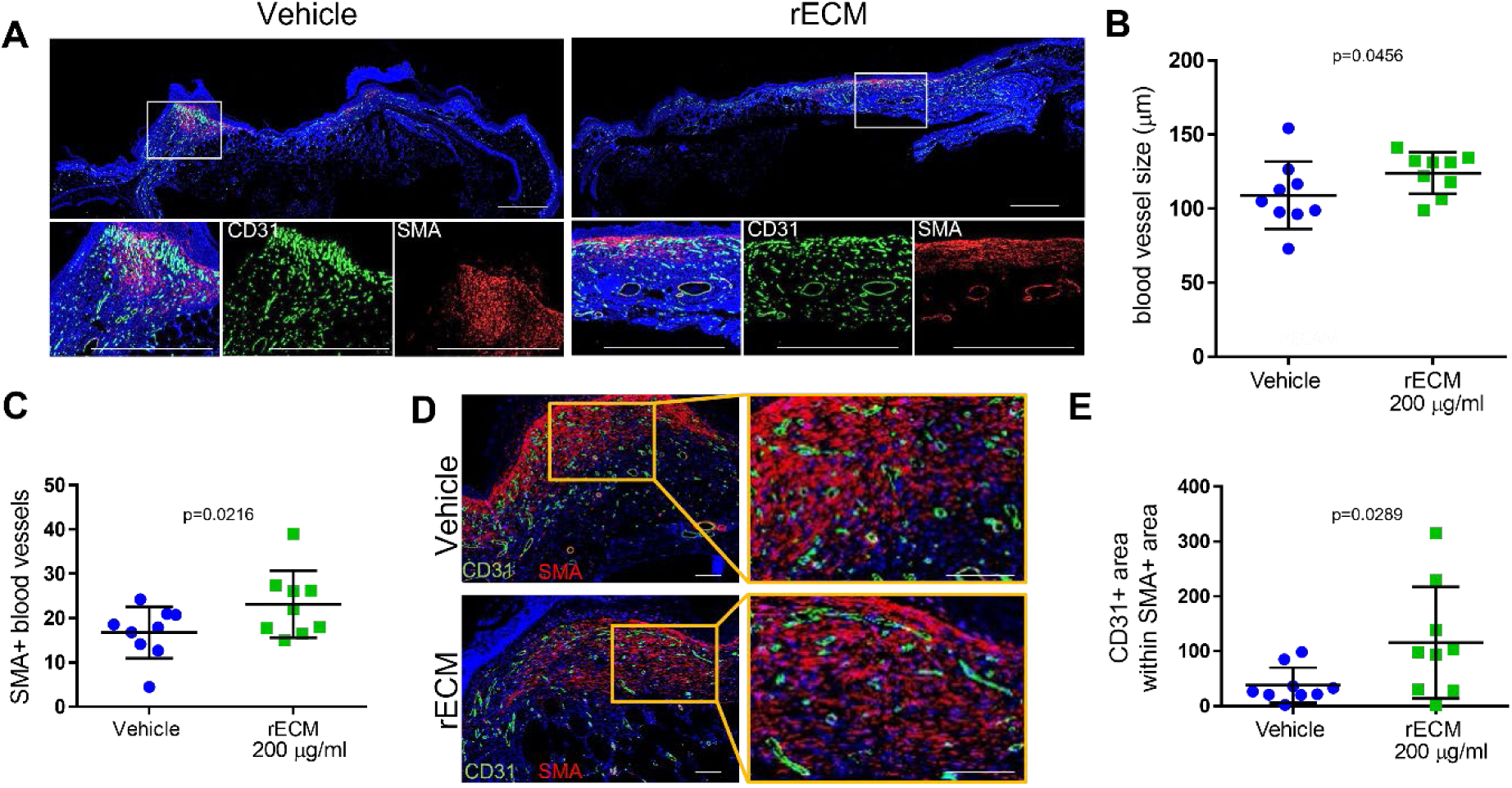
rECM accelerated blood vessel remodeling and vascular maturation *in vivo*. **Panel A:** Photographs illustrating CD31 and smooth muscle actin (SMA) staining of Day 11 skin wounds receiving Vehicle or 200 µg/ml rECM (scale bar = 1000 µm). **Panel B:** Measurements of the average size of CD31-positive blood vessels within Day 11 skin wounds receiving Vehicle or 200 µg/ml rECM. **Panel C:** As in B, but relative number of SMA-positive blood vessels within wounds. **Panel D:** As in B, but CD31 positive area within the positive smooth muscle actin (SMA) staining of myofibroblasts (scale bar = 100 µm). **Panel E:** Measurements of the average density of CD31 positive cells within myofibroblast-dense area. *<u>Statistics:</u>*All data panels: paired t-test; central bar represents mean, and error bars represent standard deviation (n = 9 mice).

To gain additional insight into the contribution of rECM to the initial stages of new blood vessel growth, we tested rECM in a defined 3 dimensional (3D) model of angiogenic sprout initiation, where angiogenic endothelial cells are induced to undergo invasion into collagen matrices.(54, 55) Collagen matrices received increasing doses of rECM, and human endothelial cells were placed on the surface prior to incubation for 24 hrs. For vehicle and rECM doses at 15.6, 31.25 and 62.5 μg/ml, robust sprouting occurred rapidly over 24 hrs (**Fig6A**). No significant differences were observed in the number of structures that invaded (**Fig6B**), but closer examination of invasion distance (the distance between the monolayer and deepest point of cell invasion) revealed that the rECM accelerated the rate of invasion (**Fig6C**). Accelerated invasion with rECM treatment also coincided with longer filopodia (**Fig6D**). At rECM doses 125 μg/ml and above, the endothelial cells unexpectedly promoted complete contraction of collagen matrices (**Fig6E,F**). Nanoindentation testing confirmed that cell-induced contraction was not caused by rECM-induced weakening of the collagen matrix (**Fig6G**). Surprisingly, high concentrations of rECM appeared to modestly increase stiffness, but under the conditions of the measurements, statistical significance was only observed with 125 μg/ml and 500 μg/ml rECM. Endothelial to mesenchymal transition (EndMT) to a myofibroblastoid phenotype (56) was considered as an explanation for the contraction phenomena as myofibroblasts are key participants in wound contraction, (57) but no increases in the expression of EndMT markers were observed in response to rECM (FigS3). Collectively, these data indicate that while rECM does not alter the ability of endothelial cells to initiate angiogenic sprouting, rECM does accelerate the process by promoting faster migration and longer filopodia formation. At higher doses, the rECM also participates in stimulating contraction of collagen matrices, which is consistent with acceleration of granulation tissue remodeling.

**Figure 6.**
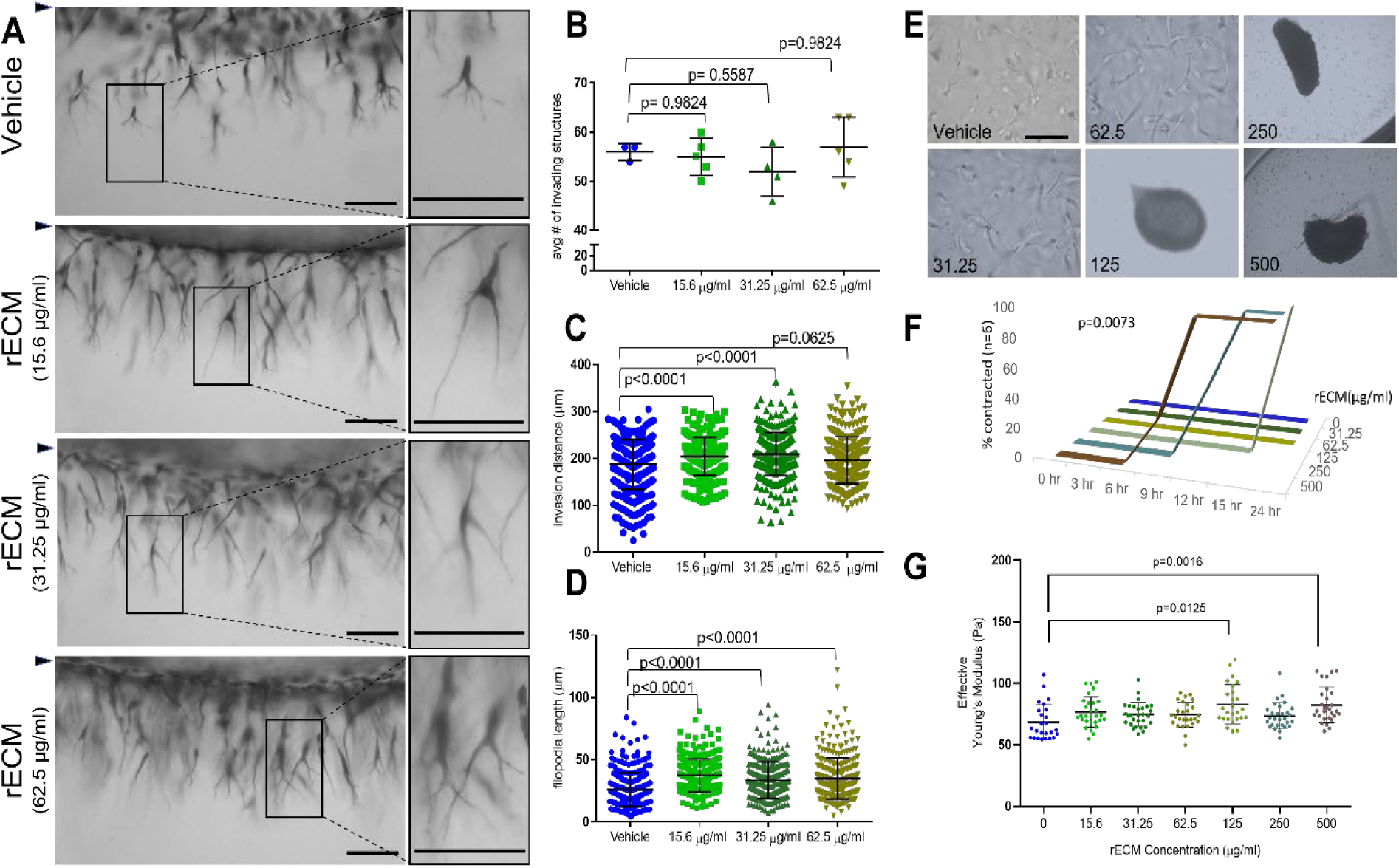
rECM modulates angiogenic sprouting *in vitro*. **Panel A:** Photographs representing human endothelial cell invasion responses in collagen matrices. A side view is shown with cells invading beneath the original monolayer (indicated by arrowhead). Magnified images of filopodia are presented on the right (scale bars *=* 100 µm). **Panel B:** Quantification of invasion density. **Panel C:** Quantification of invasion distance defined as the average distance migrated from the monolayer. **Panel D:** Quantification of filopodia length, defined as the average length of filopodia extending from the cell body. **Panel E:** Photographs of endothelial cell monolayers from above (scale bar *=* 100 µm). **Panel F:** Frequency of contraction incidence over time in culture (n = 6/group). **Panel G:** Moduli of collagen matrices containing indicated doses of rECM. *<u>Statistics</u>*: Panels B and C: Data represent the average number of invading cells per standardized field (0.25 mm^2^) and average invasion distance, n = 3-5 independent wells; central bar represents mean, and error bars represent standard deviation. Data were analyzed by one-way ANOVA. For invasion distance, n≥ 245 structures were measured. Panel D: At least 300 filopodia were measured per treatment group from 3 independent wells; central bar represents mean, and error bars represent standard deviation. Data were analyzed by one-way ANOVA. Panel G: The average Effective Young’s Modulus for each gel (n = 3 per condition). Data were analyzed by one-way ANOVA with a Dunnett’s post-hoc test, comparing each condition to the Vehicle control (0 µg/ml rECM).

## Discussion

Herein we describe a solubilized ECM preparation purified from human iPSC-derived MSCs with the capacity to improve several parameters of dermal wound healing in leptin receptor null (*db/db*) mice, a standard rodent model of type 2 diabetes. The *db/db* mice were chosen for this study because they develop obesity, hyperglycemia, insulin resistance and exhibit impaired re-epithelialization and keratinocyte migration resulting in delayed closure of full-thickness excisional wounds (40, 41).

When solubilized rECM was combined with type I collagen and applied to 8 mm full-thickness dermal defects, it significantly enhanced wound contraction between days 7 and 13 post-injury (**Fig1B, 2A**), temporally coincident with the granulation phase (**Fig 2E**). Granulation tissue accumulates rapidly following the inflammatory and hemostatic phases, driving wound contraction, neovascularization, and immunomodulatory processes. (43–45) The observation that lower doses of rECM result in thicker granulation tissue at day 11 but higher doses result in a thinner layer of granulation tissue at the same time point suggests that rECM dose-dependently accelerates both the generation and remodeling of granulation tissue (**Fig3D**). This interpretation is further supported by accelerated wound closure with rECM (**Fig1B, Fig2A**) and reduced proportion of granulation tissue at the wound site at high rECM doses (**Fig3E**). Granulation tissue is inherently transient and expedited progression through this phase by the action of rECM is expected to increase the rate and likelihood of effective repair. The mechanism by which rECM drives granulation is currently unclear but of the 20 plus matrisomal proteins present in rECM, (24) one of the most abundant proteins, collagen VI, serves as a key regulator of granulation tissue matrix formation (32) and therefore is a primary mechanistic candidate for rECM-driven healing.

The role of wound contraction and the synthesis of granulation tissue is critical for wound stabilization at the initial stages of repair, but maturation of vasculature, formation of skin appendages and reinnervation are all essential for the completion of functional skin healing (**Fig2E**). (42, 43, 45) Quantitative histological analyses were performed at day 11 because the majority of observed rECM-induced healing occurred at this time, and prior studies predicted granulation, re-vascularization, and re-epithelialization should be observable at this timepoint. (42) Spatial sequencing identified several gene signatures indicative of processes expected during wound healing (immune processes, cytokine responses, epithelial and keratinocyte signatures, angiogenesis) but no unexpected or deleterious rECM-driven phenomena (rejection, scarring, infection, neoplasia) were observed (**FigS2, TableS1**).

Neuroregulation is an underestimated process within the context of skin healing, but it is becoming increasingly apparent that the presence of appropriate neuroregulatory cues participate strongly in regulation of wound repair, angiogenesis and proliferation of resident progenitors. (60, 61) For example, peripheral glia have been shown to play a key role in wound contraction and myofibroblast proliferation, (62) sensory peptides substance P and calcitonin gene regulatory peptide accelerate skin healing through mechanisms involving angiogenic and vasodilatory effects, (63) and appropriate levels of innervation can reduce the probability of scarring in favor of re-epithelialization. (64) In this study, rECM was observed to dose-dependently increase the frequency of structures that resembled newly-formed peripheral nerve clusters (**Fig4A**). When spatial sequencing was performed on sections harboring these structures, their expression signatures corresponded strongly to Schwann cells and peripheral nerves (**Fig4C**) with prominent transcription of key Schwann cell and myelination markers such as myelin protein zero (*Mpz*), periaxin (*Prx*), myelin basic protein (Mbp) and proteolipid protein-1 (*Plp1*), and also glial-associated transcripts such as S100b and cytokine receptor-like factor-1 (*Crlf-1*).

Parfejefs et al. (62) described a population of skin-resident glial cells present in injury-activated peripheral nerve clusters that promote wound healing through paracrine modulation of transforming growth factor-β (TGF-β) signaling. In these studies, depletion of the cells perturbed skin healing through inhibition of wound closure, epithelial cell proliferation and myofibroblast activity. Parfejefs *et al.* stratified the regenerative glia into MBP or p75 positive variants, representing a differentiated functional form and a more proliferative de-differentiated form respectively. Our data demonstrate the presence of both forms of glial variants (**Fig4C,D)** in rECM-treated skin, providing one potential explanation for the acceleration of wound contraction and the granulation phase. The mechanism underpinning the enhanced development of peripheral nerve structures in response to rECM is currently unclear, but type VI and type XII collagens, both major components of rECM, promote peripheral nerve regeneration (58) and facilitate axon regrowth through provision of an attachment substrate (34), respectively. The increased prevalence of peripheral nerves in rECM also offers the clear potential for enhanced functionality, such as improved sensitivity to thermal, sensory and traumatic stimuli (65) but also raise concerns related to the potential for trauma-induced neuropathic pain that would need to be addressed in future studies. (66) It should be noted here, however, that qualitative assessment of pain responses throughout the studies did not indicate rECM-induced discomfort.

Angiogenesis promotes effective skin re-epithelialization but when it is excessive, prolonged, or structurally abnormal, it can promote fibrosis and scarring (67, 68). The maturity of angiogenic vessels strongly influences wound healing outcomes with more mature, stable, well-perfused vessels favoring efficient healing. (69) While blood vessels at various stages of maturation were visible in all treatment groups, rECM treatment resulted in larger and more regular-sized CD31-expressing blood vessels (**Fig5B**) that had a greater tendency to colocalize with smooth muscle actin-positive periendothelial cells (**Fig5C,E**). Collectively, these features are associated with advanced blood vessel maturity. (70, 69) In agreement with *in vivo* observations, more rapid endothelial sprouting responses in 3D culture experiments was observed (**Fig6A-D**). While rECM did not increase the density of invading structures, rECM dose-dependently accelerated the rate of invasion that coincided with longer filopodia. Potential explanations include the slight increase in stiffness of collagen matrices, which agrees with prior studies demonstrating that cross-linking or increasing density of collagen matrices enhanced angiogenic responses. (71–73) Sprouting is also bolstered by acquisition of an endothelial to mesenchymal transition (EndMT) phenotype in endothelial cells; (74, 75) however, we did not observe changes in EndMT marker expression with rECM treatment (**FigS3**). Surprisingly, higher rECM doses promoted complete contraction of collagen matrices that could not be explained by decreased stiffness or acquisition of an EndMT or myofibroblastoid phenotype. (56, 57, 77, 78, 79)

Measurement of wound area by digital imaging and/or calipers provided a convenient means to non-invasively assess healing in real-time, but it was subject to variation that arose from differences in methodology and the size, appearance and extent of scabbing. At least three observers blinded to the conditions of the assay significantly reduced analysis variation, but this did not fully address complications related to scab size that did not necessarily correlate with the degree of underlying healing. The multi-parameter histological assay of Van der Vyer *et al.,* (42) addresses many of the complexities of skin healing through the development of a scoring system that collectively quantifies key histological signs of healing including re-epithelialization, epidermal thickness, keratinization, granulation tissue thickness, skin appendage development, and scar elevation index (**Fig2D,E**, **Table 1**). We found this protocol to possess high fidelity, especially when slight adjustments were made to scoring criteria to suit the experimental conditions (**Table 1**). Outcome was significantly improved by careful preparation of the specimens to maintain all epidermal and hypodermal aspects of the skin morphology, standardized tissue orientation and utilization of high-resolution slide scanners with artificial intelligence-enabled image analysis software.

Leptin receptor null (*db*/*db*) mice are a powerful model for diabetic dermal wound healing in that they recapitulate many key defects (delayed closure, impaired re-epithelialization). Compared to wild-type mice, wounds heal much more slowly, with minimal early contraction, prominent wound expansion and greater reliance on slow re-epithelialization and granulation tissue formation. (40, 41) However, the *db*/*db* mice and rodents in general have limitations as models of diabetic wound healing. Leptin receptor mutations are not a common cause of type 2 diabetes, and the overrepresentation of the role of leptin signaling in this model may result in limited capacity to mimic elements of the disease in humans. (76) Furthermore, murine dorsal skin has loose attachment and a prominent panniculus carnosus that drives closure by contraction, whereas human skin heals mainly by granulation tissue formation and re-epithelialization. Even though contraction is relatively impaired in *db*/*db* mice, they still possess this muscle layer and loose skin, so wound mechanics and scar formation differ fundamentally from human foot or leg ulcers. (77) Interestingly, newly formed wounds on *db*/*db* mice were immediately enlarged (as compared to the biopsy punch), suggesting skin laxity. Sullivan *et al.* also reported this observation in *db*/*db* mice which did not occur in heterozygous counterparts (41). In this study, when rECM was applied to these defects, immediate wound expansion did not occur. The observation that this effect occurred within seconds suggests that rECM provides an immediate biomechanical contribution that counteracts skin laxity and restores some degree of stability. Dermal myofibroblasts have the capacity to respond very rapidly to contractile stimuli to contract collagen matrices within minutes. (78, 79) Periostin is a major component of rECM, (24) known to rapidly stimulate contractile responses in myofibroblasts, (80) and therefore could serve as a key functional mediator in rECM-driven skin defect closure. (81) The rapid nature of these contractile responses could be particularly useful in trauma settings where wound closure and rapid hemostasis are key determinants of patient outcome, but confirmation that these observations occur in tissue that more closely recapitulates human skin is needed. (86)

Together, these findings demonstrate that solubilized rECM derived from human iPSC-derived MSCs accelerates closure of diabetic skin defects in *db*/*db* mice by accelerating wound closure, expediting the granulation phase, promoting blood vessel maturation and augmenting peripheral nerve regeneration. By recapitulating key structural and instructive features of native matrix, this acellular biomaterial offers a practical, cell-free strategy to enhance functional wound repair in challenging diabetic settings and warrants further investigation as a translational therapy for chronic non-healing wounds.

## Materials and Methods

### Cell Culture

hMSCs were cultured as previously described. (25) Briefly, hMSCs were grown in α-MEM supplemented with an additional 2 mM L-glutamine (4 mM final), 20% FBS (fetal bovine serum), 100 U/ml Penicillin, and 100 µg/ml Streptomycin in 150 cm^2^ culture dishes. Cells were grown in a humidified 37°C incubator with 5% CO_2_, with media changes every second day until reaching 80% confluency, at which point they were switched to α-MEM, 2 mM L-glutamine (4 mM final), 20% FBS, 100 U/ml Penicillin, 100 µg/ml Streptomycin, 1% L-Glutamine, 5 mM β-Glycerophosphate, 50 µg/ml Ascorbic Acid, 10 µM GW9662 and fed every second day for 10 days. Human umbilical vein endothelial cells (ECs) were cultured at passages 3–6 (Lonza, NJ) in 75 cm^2^ flasks (Corning) precoated with 1 mg/ml sterile gelatin. Growth medium was previously described in detail (55) and consisted of M199, 10% FBS, 0.4 mg/ml bovine hypothalamic extract (Pel-Freeze Biologicals, AR), 100 µg/ml heparin, and penicillin, streptomycin and gentamicin.

### Preparation of Collagen Matrices

Collagen matrices were prepared as previously described. (55) Briefly, type 1 collagen was extracted from rat tails (Pel-Freeze Biologicals, AR) by incubation in sterile 0.1% acetic acid before lyophilization and resuspension at 7.1 mg/ml in 0.1% acetic acid. Collagen matrices (2.5 mg/ml) were prepared by combining collagen with 10X M199 (Gibco), 5N NaOH, and M199 (Gibco). (55)

### Preparation of Regenerative Extracellular Matrix (rECM)

The regenerative Extracellular Matrix (rECM) was prepared as previously described. (25) Briefly, osteogenic monolayers were frozen at –80°C for at least 15 hrs before suspension in 1X PBS supplemented with 0.1% (v/v) Triton X-100, 1 mM MgCl_2_, and 1 U/ml DNAse 1 for 2 hrs, at which time, 0.1% (v/v) trypsin was then added for 15 hrs. To purify the rECM, several ice-cold water and chloroform washes were performed before a final rinse with acetone. After air drying for at least 4 hrs, the rECM was weighed and resuspended at 1 mg/ml in 0.1M acetic acid. Protein concentrations were confirmed by measuring absorbance at 280 nm.

### In vitro Angiogenesis Assays

Confluent ECs were trypsinized and seeded at 30,000 cells per well on 2.5 mg/ml collagen matrices prepared as described (55) containing 1 μM S1P (Sphingosine 1-phosphate) (Avanti Polar Lipids, AL), along with indicated concentrations of rECM. Collagen (25 µl) was added to 96 well half area plates (Costar, NY) and equilibrated for 45 min. All media included M199 supplemented with ascorbic acid, RSII and 40 ng/ml VEGF and bFGF.(55) Humidified cultures were allowed to invade for 20–24 hrs, unless otherwise indicated, at 37°C with 5% CO_2_ before fixation in PBS containing 3% glutaraldehyde (Sigma, MO) or 4% paraformaldehyde (Electron Microscopy Sciences, PA).

After overnight fixation, gels were stained with 0.1% toluidine blue in 30% methanol. For invasion density measurements, a minimum of three fields were quantified with QCapture Pro acquisition software. For invasion distance, at least 50 structures from each treatment group were included in the analysis, where the distance migrated from the monolayer was recorded using side view images. Image-Pro PLUS (MediaCybernetics, MD) software was used to quantify invasion distance and peripheral process extension. At least 30 cells per treatment group were analyzed for the number of peripheral processes using toluidine blue stained cultures. To assess contraction of cultures, time courses were performed, and collagen matrix diameter within each well was recorded. Contraction was detected by collagen matrices pulling away from the side of the well and eventually condensing into condensed cell aggregates, indicative of 100% contraction.

### Nanoindentation

An Optics11 Life Pavone Nanoindenter was used to assess the effects on the Young’s Modulus of the collagen matrices when rECM was incorporated at increasing amounts. A probe (stiffness = 0.025 N/m, tip radius = 25 um) was used to indent the surface of the matrix in a 3×3 grid. The Effective Young’s Modulus was calculated for each individual point and averaged per well. The average Effective Young’s Modulus for each matrix (n=3 per condition) was used in a One-Way ANOVA with a Dunnett’s post-hoc test, comparing each condition to the Vehicle.

### Mouse Wound Assays

All animal protocols and procedures were performed as defined in the Animal Use Protocol 2024-0292 approved by Texas A&M University. Twenty (10 male and 10 female) 12-week-old leptin receptor-deficient (*db/db*) diabetic mice were shaved one day prior to wounding with depilatory cream. The rECM was incorporated into 2.5 mg/ml collagen matrices at 0 (0.1M acetic acid Vehicle) or 100 µg/ml rECM concentrations. Mice received subcutaneous analgesic prior to being anesthetized. A 8 mm sterile punch was used to place two full thickness skin wounds in the cleared areas. Wounds were filled with collagen matrices containing either Vehicle or rECM, which solidified upon contact with the wound and formed a plug. Wounds were covered with liquid Band-Aid, and wounds were photographed thereafter to monitor wound closure until tissue collection on day 20. Photographs were taken of each set of wounds on days 0-3, 5-10 and 13-20 and analyzed by 3 individual observers using Image J to calculate the area of the wound. The average wound area from 3 observers, from all mice for each day for each treatment, Vehicle or rECM were averaged and a 2-way ANOVA performed. SEM bars on graphs represent the variance in the mice wounds per side per day. Additionally, a paired, Student’s t-test was used to calculate the defect area significance of the Vehicle versus rECM treated wounds each day, with error bars representing standard deviation.

In a separate histological study, 12-week-old leptin receptor deficient (*db/db*) diabetic mice received two 8 mm full thickness skin wounds as described above and were divided into 3 groups. Each wound received Vehicle (0.1M acetic acid) and either 50, 100, or 200 µg/ml rECM. Each group contained 5 female and 5 male mice. Mice were observed and monitored daily, and caliper wound measurements and photographs were collected at days 5, 8, and 11 to assess wound closure and healing. Animals were euthanized prior to tissue collection 11 days after wounding. Tissues were fixed in 3.7% formalin in PBS overnight, and wounds were cut in half before embedding in paraffin. 4 µm sections were cut, mounted onto charged glass slides. H&E and Masson’s Trichrome stains were performed on serial sections at the VMBS Core Histology Lab, Texas A&M University.

### Histological Imaging and Analysis

Using a VS120 Slide Scanner at the IMIL (Integrated Microscopy and Imaging Laboratory) at Texas A&M University, pyramidal files were collected and analyzed using Qu-Path-0.5.1. The methods from van de Vyver et al. (42) were adapted to assign a healing index score on a scale of 0 (unhealed and open) to 12 (completely healed without excessive scarring) to Vehicle and rECM treated wounds using Masson’s Trichrome stained sections. Six parameters were scored by at least 4 observers. (42) Due to slight variations in the experimental healing, criteria were adapted as necessary for the current study, indicated below as “Adapted Criteria” (**Table 1**).

The 6 parameters above contributed to the overall wound healing score. Additional analyses using Qu-Path included granulation tissue area, number of appendages in the wound area, and the number, area and clusters of Schwann cells present.

### Spatial Sequencing

Two formalin fixed, paraffin (FFPE) embedded rECM wound samples were submitted for spatial sequencing to the Institute for Genome Sciences and Society Molecular Genomics Core Texas A&M University. After passing the quality control, a specific 6.5 mm x 6.5 mm area was chosen for sequencing on the 10x Genomics Visium platform. The 10x Genomics Space Ranger pipeline and Loupe Browser 8 were used to analyze the spatial seq data. Unidentified structures in Masson’s Trichrome stained sections were identified through analysis of differentially expressed genes within the “spots” localized to these structures using GOnet pathway software (https://tools.dice-database.org/GOnet/), and bubble plots generated with the use of SRPLOT (https://www.bioinformatics.com.cn/plot_basic_gopathway_enrichment_bubbleplot_081_en). Using Spot Deconvolution in Loupe Browser, tissue sections were annotated with “k” number of topics, (maximum 7 for Sample 1 and 8 for Sample 2) based on their gene expression profile with default settings. Each k topic represents cell types or mixtures that are designated according to the gene expression signatures that appear simultaneously in the tissue spots. Evaluating the gene expression using GOnet pathway software (https://tools.dice-database.org/GOnet/) we were able to classify topics (**Table S1**).

### Immunofluorescence

Paraffin sections (4 µm thick) were adhered to charged glass slides and deparaffinized with xylene and hydrated through decreasing ethanol washes. Antigen retrieval was done in citrate buffer (10 mM citric acid, 0.05% Tween 20, pH 6.0) with 15 psi at 121°C. Sections were blocked with 5% normal goat serum in 1X PBS (blocking buffer) for 1 hr before addition of primary antibodies. Primary antibodies were diluted in blocking buffer at indicated concentrations, applied to sections and incubated at 4°C overnight. Sections were washed 3 times for 5 minutes each in 1X PBS. Goat anti-rat or goat anti-rabbit Alexa Fluro 488 or 594 conjugated secondary antibodies diluted 1:300 in blocking buffer were added to sections for 1 hr at room temperature, protected from light. Sections were washed 3 times for 5 minutes each in 1X PBS. Samples were stained with 10.9 µM DAPI for 5 minutes and rinsed in 1X PBS. Slides were incubated with True Black (Cat# 23007, Biotium, CA) as per manufacturer’s suggestions to reduce autofluorescence of red blood cells. Sections were rinsed in 1X PBS before mounting with Fluorgel (Electron Microscopy Sciences, PA) and then imaged on an Olympus Fluoview 3000 confocal microscope with FV 31S-SW acquisition software. Antibodies used are listed in **Table S3**.

### Immunofluorescence Quantification

Fluorescent signal was quantified using Fiji. Briefly, .oir files were uploaded into Fiji and ROIs of wound area were identified and measured. Thresholds were set identical for all images within a group, and signal area was measured using the analyze particles tool. Data are displayed as signal area normalized to wound area, unless otherwise indicated.

### RNA Extraction and qPCR

Cells were allowed to invade for 3 hrs before digestion in RLT Lysis Buffer, and RNA was extracted with Qiagen’s RNeasy Kit. cDNA was made using Invitrogen’s SuperScript III cDNA synthesis kit with 0.5 µg RNA as a starting template and oligo(dT)_20_ as per manufacturer’s protocol. qPCR was performed using PowerUp SYBR Green Master mix (Thermo Fisher, MA), 0.5 µmol/L forward and reverse primers and 1-µl diluted cDNA template, on a StepOnePlus Real-time System (ABI, MA) for data analysis as 2^−ΔΔCT^. RPLP0 and GAPDH were used as the housekeeping gene. Primer sequences are listed in **Table S4**.

### Statistical Analysis

GraphPad Prism version 6.07 for Windows (GraphPad Software, La Jolla, CA, www.graphpad.com) was used for post hoc statistical analyses. Raw data values were analyzed for normality and equal variation before the application of parametric or nonparametric statistical tests. Individual tests applied are indicated in each figure, along with exact *P* values. Error values represent SD or SE as indicated on each figure.

## Acknowledgments

The authors acknowledge the assistance of the Integrated Microscopy and Imaging Laboratory at the Texas A&M University Naresh K. Vashisht College of Medicine. RRID:SCR_021637. We thank the Texas A&M VMBS Core Histology Laboratory for their assistance with tissue processing and staining for this project.

## Funding and COI statement

A pilot grant to KJB and CAG from the Texas A&M Naresh K. Vashisht College of Medicine Seedling Grant Program. R01-DE032031 (CAG) National Institute for Dental and Craniofacial Research. All authors declare no potential competing interests related to the content of this manuscript.

## Authors’ contributions

Experimental design: CG, KJB

Data collection: CAA, JB, EG, JM, SL, IJ, CS, AH, HA

Data Analysis: CAA, JB, EG, JM, SL, IK, IJ, AK, CS, AH, HA, KJB, CG

Manuscript drafting: CG, KJB, CAA

Manuscript editing and approval: all authors

## Data Availability Statement

The datasets generated and/or analysed during this study are available in the Texas Data Repository, https://urldefense.com/v3/ https://dataverse.tdl.org/dataverse/rECM_;!!KwNVnqRv!H2aVgEkUbENGjuK07TM77X3WY2ndFFYZte8u-uzyGF6ktkRdW_T0GrpzI2ogwQOcSTckPXWPwbE5FeFLF-g$)

---

**Supplemental Figure S1:**
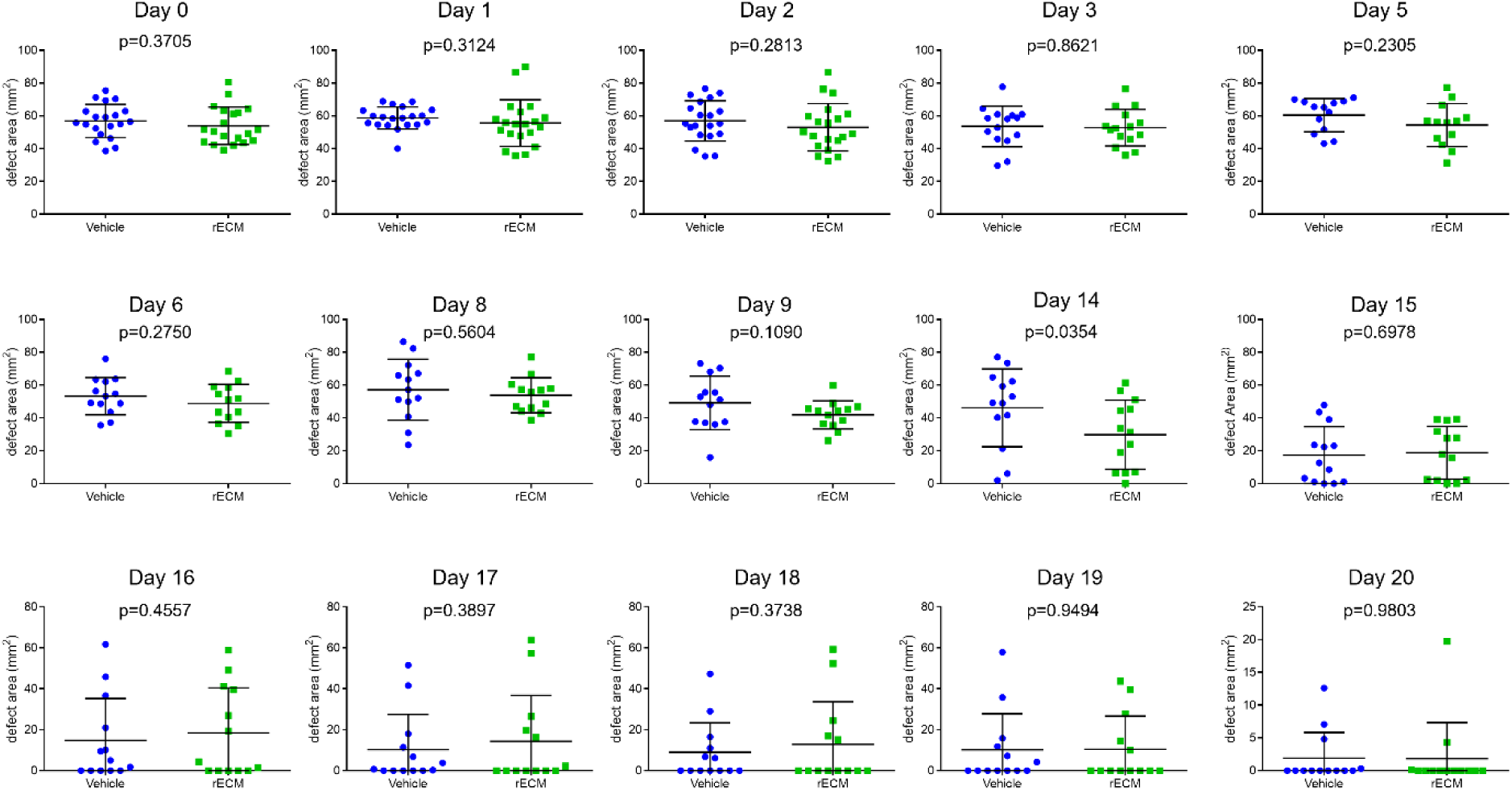
rECM accelerated dermal wound closure. Each leptin receptor deficient mouse received two 8 mm diameter, dermal defects that received Vehicle or 100 µg/ml rECM in a collagen type I matrix (2.5 mg/ml). Defect areas are plotted over time. Defect area represents the mean of 3 independent observers’ analysis of digital images. *<u>Statistics</u>*: p-values calculated by paired Student’s t-test. (n=13-20 mice).

**Supplemental Figure S2:**
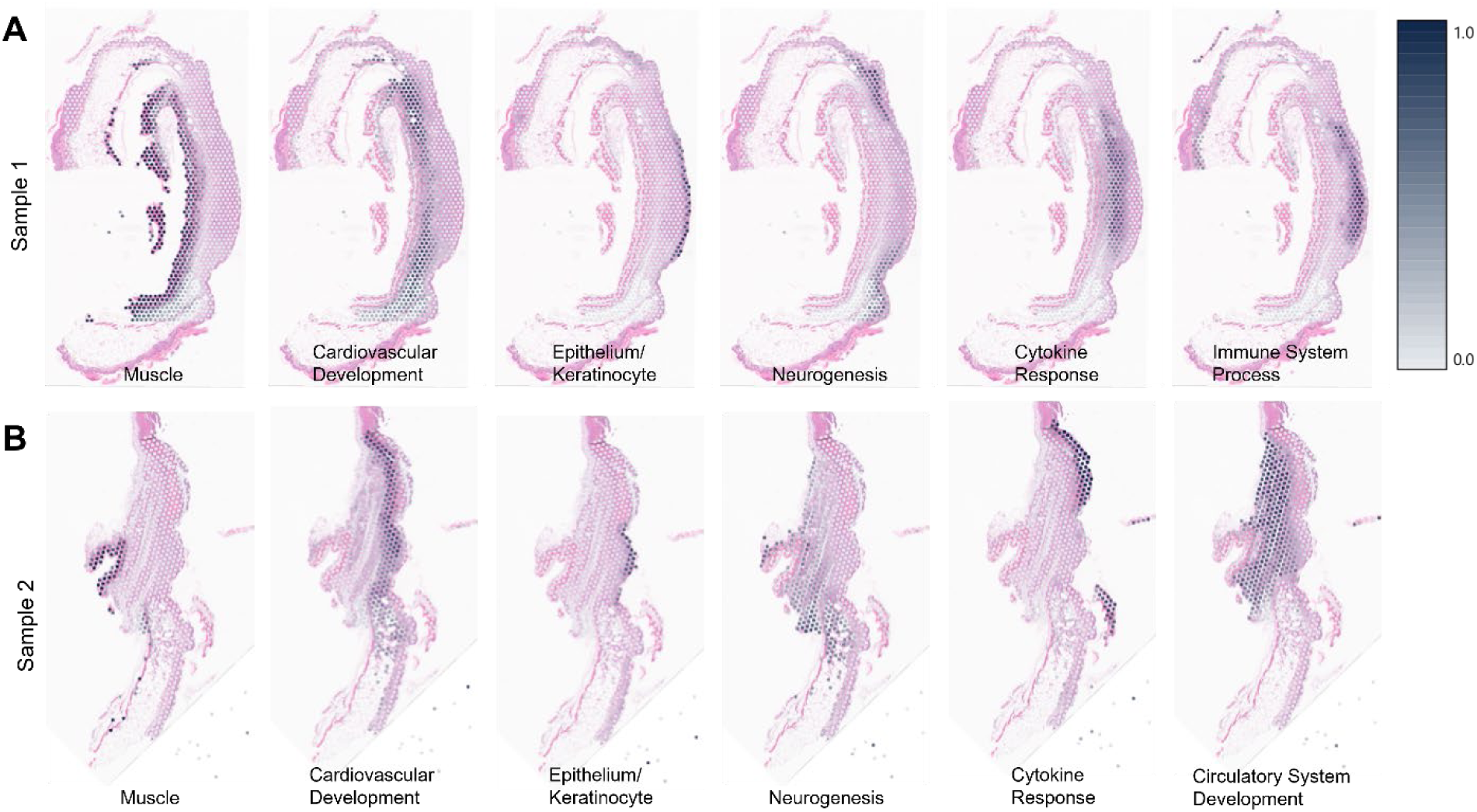
Spatial sequencing profiles in rECM-treated skin wounds. **Panels A and B:** Spatial sequencing of two independent healing skin sections with circular overlays representing individual transcriptomic datasets. Gray circles represent enrichment for transcriptomic expression of gene expression profiles indicated.

**Supplemental Figure S3:**
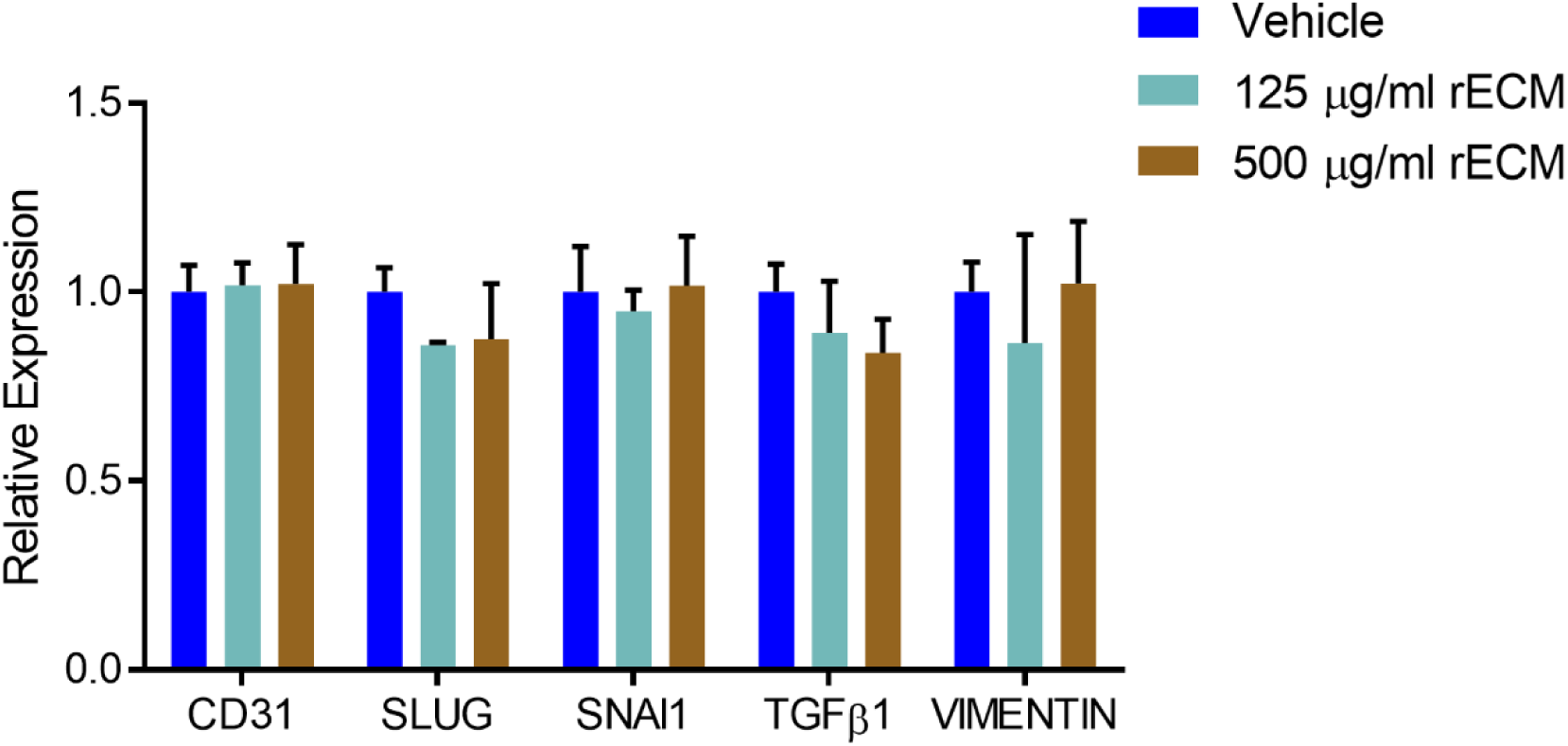
Expression of EndMT markers in endothelial cells within sprouting assays with and without rECM treatment. Relative gene expression in invading endothelial cells (3 hrs) determined using qPCR. Data shown are representative means, and error bars represent standard deviation from two independent experiments.

**Table 1: Adjustments to scoring criteria of histological wound healing.**

**Table S1: GOnet analysis of clusters identified in spatial sequencing.**

**Table S2: Schwann Cell Cluster Expression**

**Table S3: Antibodies**

**Table S4: Primer Sequences**

## Notes

### Competing Interest Statement

The authors have declared no competing interest.

